# *DLX5/6* GABAergic expression affects social vocalization: implications for human evolution

**DOI:** 10.1101/2020.07.24.218065

**Authors:** Giovanni Levi, Camille de Lombares, Cristina Giuliani, Vincenzo Iannuzzi, Rym Aouci, Paolo Garagnani, Claudio Franceschi, Dominique Grimaud-Hervé, Nicolas Narboux-Nême

**Affiliations:** Physiologie Moléculaire et Adaptation, CNRS UMR7221, Département AVIV, Muséum National d’Histoire Naturelle, Paris, France; Laboratory of Molecular Anthropology & Centre for Genome Biology, Department of Biological, Geological and Environmental Sciences, University of Bologna, Italy; Alma Mater Research Institute on Global Challenges and Climate Change, University of Bologna, Italy; Department of Experimental, Diagnostic and Specialty Medicine (DIMES), University of Bologna, Bologna, Italy; Clinical Chemistry, Department of Laboratory Medicine, Karolinska Institutet at Huddinge University Hospital, Stockholm, Sweden; Institute of Information Technologies, Mathematics and Mechanics, Lobachevsky University, Nizhniy Novgorod, Russia; Histoire Naturelle de l’Homme Préhistorique, CNRS UMR 7194, Département H&E, Muséum National d’Histoire Naturelle, Paris, France

**Keywords:** *DLX5/6* genes, ultrasonic vocalization, aging, social behaviour, Human evolution, self-domestication

## Abstract

DLX5 and DLX6 are two closely related transcription factors involved in brain development and in GABAergic differentiation. The *DLX5/6* locus is regulated by FoxP2, a gene involved in language evolution and has been associated to neurodevelopmental disorders and mental retardation. Targeted inactivation of *Dlx5/6* in mouse GABAergic neurons (*Dlx5/6^VgatCre^* mice) results in behavioural and metabolic phenotypes notably increasing lifespan by 33%.

Here, we show that *Dlx5/6^VgatCre^* mice present a hyper-vocalization and hyper-socialization phenotype. While only 7% of control mice emitted more than 700 vocalizations/10min, 30% and 56% of heterozygous or homozygous *Dlx5/6^VgatCre^* mice emitted more than 700 and up to 1400 calls/10min with a higher proportion of complex and modulated calls. Hyper-vocalizing animals were more sociable: the time spent in dynamic interactions with an unknown visitor was more than doubled compared to low-vocalizing individuals.

The characters affected by Dlx5/6 in the mouse (sociability, vocalization, skull and brain shape…) overlap those affected in the “domestication syndrome”. We therefore explored the possibility that DLX5/6 played a role in human evolution and “self-domestication” comparing *DLX5/6* genomic regions from Neanderthal and modern humans. We identify an introgressed Neanderthal haplotype (*DLX5/6*-N-Haplotype) present in 12.6% of European individuals that covers *DLX5/6* coding and regulatory sequences. The *DLX5/6*-N-Haplotype includes the binding site for GTF2I, a gene associated to Williams-Beuren syndrome, a hyper-sociability and hyper-vocalization neurodevelopmental disorder. The *DLX5/6*-N-Haplotype is significantly underrepresented in semi-supercentenarians (>105y of age), a well-established human model of healthy ageing and longevity, suggesting their involvement in the co-evolution of longevity, sociability and speech.

## INTRODUCTION

Within groups of animals, vocal calls play an essential role in directing social behaviour, in transmitting information about food availability and predators and for reproductive and maternal activities. In humans, vocal communication and speech constitute major socialization determinants; their dysfunction represents a core trait of neurodevelopmental conditions such as Autism Spectrum Disorders (ASDs). Several mouse models characterized by genetic lesions associated to human neurodevelopmental disorders have been used to study the genetic and neurological mechanisms supporting the interplay between vocalization and socialization (Caruso, et al. 2020). Mice social behaviours are accompanied by the production of ultrasonic vocalizations (USVs) (Sales 1972), which, in the case of most laboratory strains, present a frequency included between 35 and 110 kHz (Holy and Guo 2005). Mice USVs profiles vary with the type of social activity performed (e.g. courtship, aggression, fear), with the animal’s developmental stage, with their genetic background and can be altered by induced genetic modifications. Although the link between USVs profiles and social activity is still incompletely analysed, a recent study has implemented artificial intelligence algorithms to associate USVs of individual mice to specific social actions (Ey, et al. 2020; Sangiamo, et al. 2020) unveiling a functional correlation between specific USVs and social behaviours. The generation of voluntary structured vocalizations requires the coordination of different levels of cognitive and motor activities and involves a complex and diffuse neural network ranging from the prefrontal and motor cortical areas for voluntary initiation of vocalization to several forebrain and brainstem regions (Jurgens 2002; Simonyan 2014) for the control of laryngeal movements and respiration. The inputs/outputs to this network are integrated in the midbrain periaqueductal grey (PAG), the central neural region controlling vocalization in most mammalians (Jurgens 2009; Tschida, et al. 2019).

The study of developmental disorders affecting vocal communication such as stuttering, verbal dyspraxia and some types of ASDs has led to the identification of genes involved in the control of speech and/or socio/cognitive faculties (Konopka and Roberts 2016). The most intensively studied of them, the transcription factor FOXP2, is associated to inherited dyspraxia and poor control of the facial musculature. Other “language-associated” genes include for example: *SHANK1/2/3, FOXP1, GNPTAB, GNPTG, CNTNAP2*, and *NAGPA*; for most of these genes mouse models have been generated resulting almost invariably in alteration of USV and social behaviour (Wohr 2014; Caruso, et al. 2020). These animal models suggest that a certain level of conservation exists between the mechanisms controlling vocalization. Homozygous disruption of *Foxp2* in the mouse cerebral cortex results in abnormal social approach and altered USVs (Medvedeva, et al. 2019). One of the targets of Foxp2 is the *Dlx5/6* locus: the mouse line Foxp2-S321X, which does not produce Foxp2 protein, displays a more than 200-fold increased expression of *Dlx6as1* (also known as *Evf1/2* or *Shhrs*) (Vernes, et al. 2006) a long noncoding RNA generated from the *Dlx5/6* locus known to play a central role in the control of *Dlx* gene expression (Faedo, et al. 2004; Feng, et al. 2006).

*DLX5/6* are pleiotropic genes that control multiple aspects of embryonic development (Merlo, et al. 2000), including limb, craniofacial and laryngeal morphogenesis (Beverdam, et al. 2002; Depew, et al. 2002; Robledo, et al. 2002; Conte, et al. 2016) and GABAergic neuronal differentiation and function (Perera, et al. 2004; Wang, et al. 2010; Cho, et al. 2015; de Lombares, et al. 2019).

In humans, mutations in a 1.5 Mb genomic region including *DLX5/6* are associated with Split Hand/Foot Malformation type 1 (SHFM1) (Scherer, et al. 1994; Crackower, et al. 1996), a form of ectrodactyly which can be accompanied by sensorineural deafness, craniofacial malformations with or without intellectual disability (Birnbaum, et al. 2012; Rasmussen, et al. 2016). Recently a largescale transcriptomic study on post-mortem brains has suggested that the *DLX5/6* locus participates to genetic modules involved in ASDs and schizophrenia (Gandal, et al. 2018).

The general inactivation of *Dlx5/6* in the mouse results in perinatal death with a SHFM1 phenotype associated to severe craniofacial defects (Beverdam, et al. 2002; Depew, et al. 2002) while their targeted inactivation only in cephalic neural crest cells (CNCCs) results in more limited craniofacial defects (Shimizu, et al. 2018). In the brain *Dlx5/6* are selectively expressed by GABAergic neurons. Recently, by crossing *Dlx5/6^flox^* mice with *Vgat^cre^* transgenic animals in which the “*Cre-recombinase*” gene is expressed selectively by GABAergic neurons, we have generated mice in which the Dlx5/6 DNA-binding capacity is reduced (*Dlx5/6^VgatCre/+^*) or suppressed (*Dlx5/6^VgatCre^*) only in GABAergic neurons (de Lombares, et al. 2019). *Dlx5/6^VgatCre/+^* and *Dlx5/6^VgatCre^* mice present reduced anxiety, compulsivity and adiposity associated to a 33% life expectancy extension (de Lombares, et al. 2019). In this study we show that *Dlx5/6^VgatCre/+^* and *Dlx5/6^VgatCre^* mice present also different social behaviours and vocalization patterns when compared to their littermates.

It has long been noted that morphological differences between modern and archaic Homo is similar to those of domesticated species (Boas 1938). The set of characters affected by Dlx5/6 in the mouse are reminiscent of those distinguishing domesticated species from their wild counterparts (Wilkins, et al. 2014), the so-called “domestication syndrome” characterized by increased socialization and by morphological changes in brain-case shape and size, retraction of the face, and decreased tooth size (Sánchez-Villagra and van Schaik 2019). A similar set of morphological and behavioural changes also occurred in wild house mice that adapted to life in close contact to humans (Geiger, et al. 2018). The “self-domestication hypothesis” suggests that human evolution (Zanella, et al. 2019) can be better understood in the light of a domestication process leading to increased socialization and communication accompanied by progressive changes in morphological characters such as craniofacial features.

Modern humans, the sole survivor of the group of hominins, differ from other *Hominidae* by many anatomical and physiological characters and, most remarkably, for their cognitive and linguistic capacities which reflect the singularity of our brain. Comparison of modern and archaic *Homo* genomes has played a pivotal role in the quest of *Homo sapiens* evolutionary history and in defining what makes us modern humans (see for ex. (Prufer, et al. 2014; Racimo, et al. 2014; Racimo 2016; Peyregne, et al. 2017; Hajdinjak, et al. 2018).

Comparison of *Neanderthal, Denisovan* and modern human genomes has shown that interbreeding occurred between these different human groups until 45,000-65,000 years ago (Sankararaman, et al. 2014; Dannemann, et al. 2016). During later evolution, there has been strong selection against archaic introgression of Neanderthal or Denisovan sequences in the *Homo sapiens* genome while some introgressed regions were conserved and were even under positive selection. There is increasing interest to decode the biological, phenotypic and medical consequences of Neanderthal legacy in the modern human genome (Simonti, et al. 2016; Dolgova and Lao 2018; Rotival and Quintana-Murci 2020).

To understand the coevolution of human traits, a particular attention has been given to pleiotropic genes, which can simultaneously control the development of numerous characters. For example, in prospective studies trying to identify genes capable of affecting simultaneously craniofacial shape, osteogenesis and language abilities Boeckx and Benitez-Burraco have proposed a network of physically or functionally interconnected genes including the *RUNX*, *FOXP* and *DLX* families and their regulators (Boeckx and Benitez-Burraco 2014b, 2015).

Here we show that inactivation of *Dlx5/6* in mouse GABAergic neurons results in a hyper-vocalization phenotype and in a sharp increase of dynamic social interactions. Furthermore, we identify an introgressed Neanderthal haplotype (*DLX5/6*-N-Haplotype) present in 6.3% of Europeans alleles which covers *DLX6* and *DLX5* coding and regulatory sequences including SNPs associated to either hyper- (Williams-Beuren syndrome) or hypo- (ASDs) sociability and vocalization syndromes (Poitras, et al. 2010).

## RESULTS

### *Dlx5/6* expression in GABAergic neurons affects mouse social ultrasonic vocalizations

To analyse the social ultrasonic vocalization phenotypes (UVs) resulting from the simultaneous deletion of *Dlx5/6* homeodomains in adult GABAergic neurons, we applied the resident/intruder experimental paradigm to groups of control, *Dlx5/6^VgatCre/+^* (heterozygous deletion) and *Dlx5/6^VgatCre^* (homozygous deletion) adult female mice (Moles, et al. 2007).

In this experimental setting, the animals to be tested (resident) are isolated 3 days before being exposed to an unfamiliar group-housed control adult mouse (intruder). Although UVs are simultaneously recorded for the pair of mice (resident and intruder), previous studies have shown that the contribution of the new comer is minor in comparison to that of the occupant (Hammerschmidt, et al. 2012).

The number and types of vocalizations emitted during a 10min test period (v/10min) by 10 *Dlx5/6^VgatCre/+^*, 9 *Dlx5/6^VgatCre^* and 15 control females confronted to a control female intruder were individually recorded. Most (14/15; 93%) control mice emitted less than 700 v/10min and only one emitted 849 v/10min. In contrast, 30% (3/10) of the *Dlx5/6^VgatCre/+^* and 56% (5/9) of the *Dlx5/6^VgatCre^* mice presented a unique vocalization profile emitting more than 700 v/10min, with two heterozygous and four homozygous emitting more than 1000 v/10min and up to 1400 v/10min (Figure 1 A, B).

**Figure 1:**
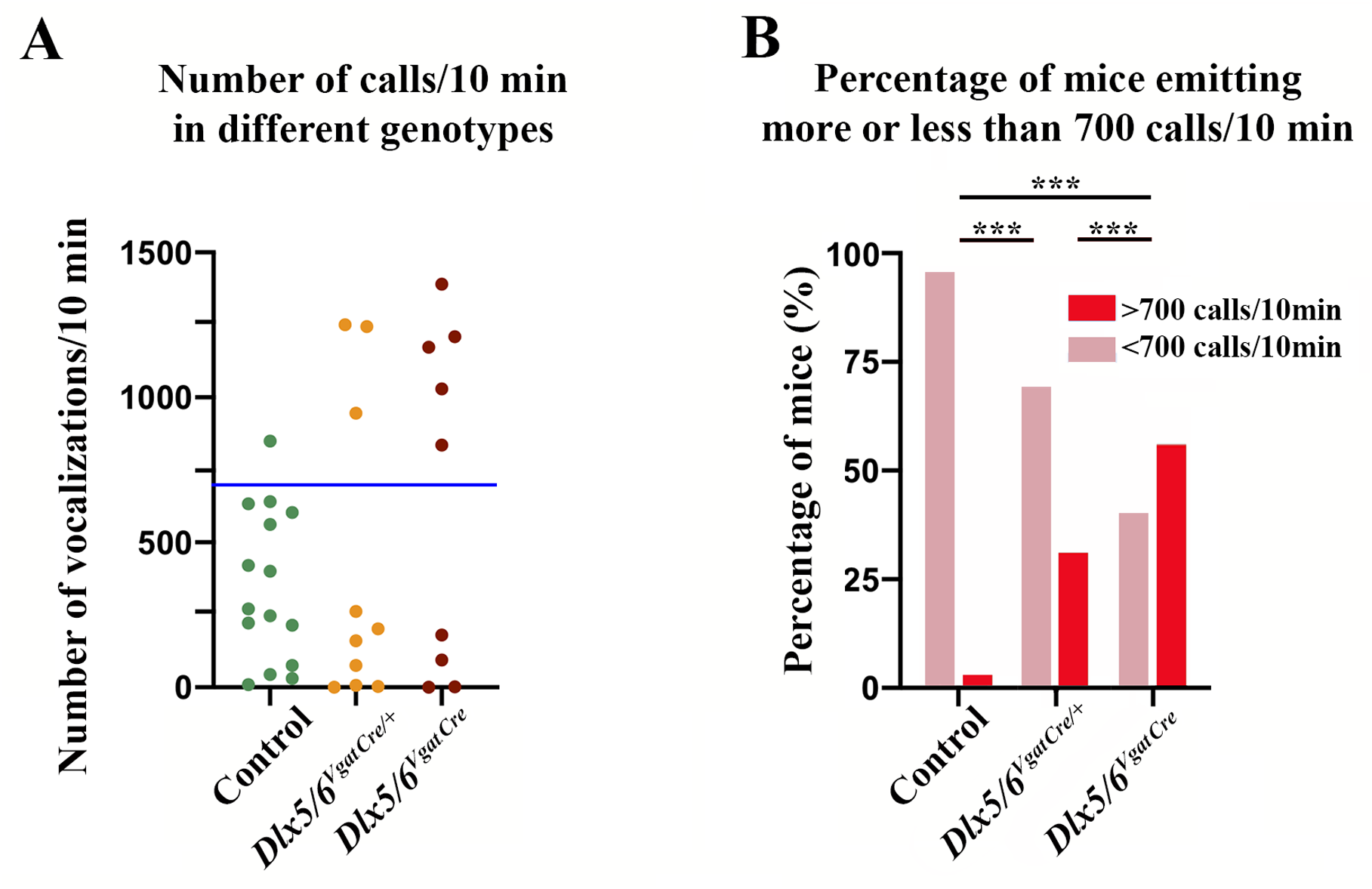
Number of calls/10 min emitted by *Dlx5/6^VgatCre/+^* and *Dlx5/6^VgatCre^* mice. Number of calls emitted by of *Dlx5/6^VgatCre/+^* (n=10), *Dlx5/6^VgatCre^* (n=9) and control (n=15) six months old female mice confronted to an intruder genetically normal female mouse of the same age over a period of 10 minutes. The number and types of vocalization were quantified. A) 93% of control mice emitted less than 700 vocalizations during the 10 minutes tests, only one outlier emitted 849 calls. In contrast, 30% of *Vgat^ΔDlx5-6/+^* (3/10) and 56% of *Vgat^ΔDIx5-6^* (5/9) mice emitted between 850 and 1400 calls/10 min. B) Percentage of mice emitting less (pink) or more (red) than 700 calls/ 10 min. The difference between each genotype is highly significant (***= p<0.001, Chi-square test).

This observation demonstrates the presence of an incompletely penetrant vocalization phenotype associated with the reduction of the level of expression of Dlx5/6 in GABAergic neurons characterized by an alleledosage-dependent increase in the number of calls per minute.

Figure 2 (A-H) presents some illustrative examples of vocalization profiles recorded from normal and mutant resident mice. Control animals (Fig. 2 A-D) and low-vocalizing mutants (Fig. 2 G, H) emitted bursts of vocal signals separated by silent intervals or did not emit any call for long periods of time (Fig. 2H). In contrast, the high-vocalizing *Dlx5/6^VgatCre^* mice emitted uninterrupted series of calls extending for many seconds without any detectable interruption or pause (Fig. 2 E, F), this pattern of vocalization being never observed in control animals.

**Figure 2:**
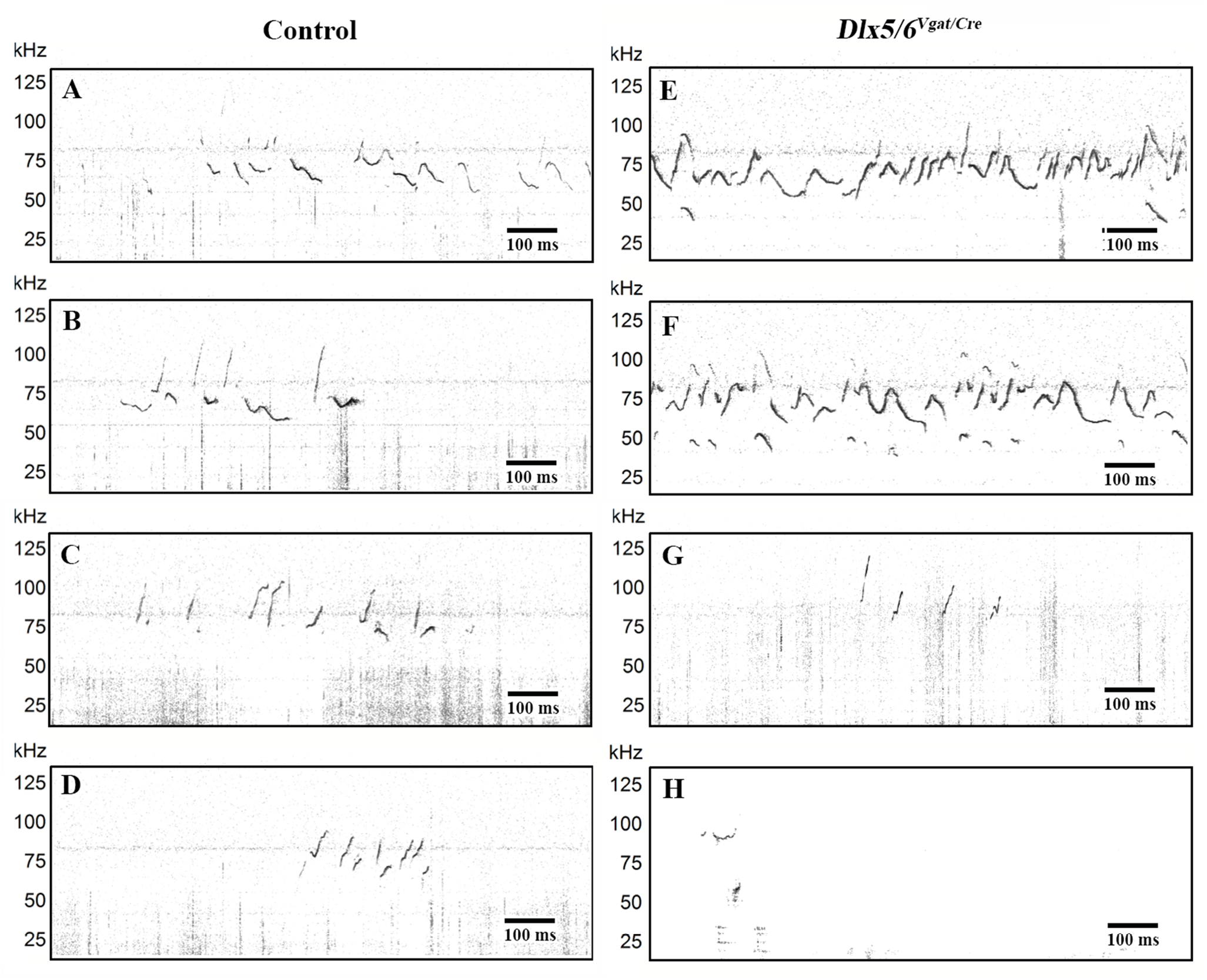
Spectrograms of control and *Dlx5/6^VgatCre^* mice. Representative spectrograms from 4 controls (A-D) and 4 *Dlx5/6^VgatCre^* (E-H) mice. In each group of animals some individuals (e.g. D, H) emitted very few vocalizations while others emitted sequences of calls separated by silent intervals (A-C, G). Hyper-vocalizing *Dlx5/6^VgatCre^* mice emitted long, uninterrupted, sequences of calls (E, F) which were never observed in controls or low-vocalizing animals.

The total time spent vocalizing by mice emitting a high number of vocalizations was about 4 times longer than that of low-vocalizing littermates (Figure 3A). We categorized the types of vocalization emitted in five major groups short, simple, complex, frequency jumps and unstructured as previously described (Ey, et al. 2013). The analysis of the specific types of calls emitted by each animal showed that high-vocalizing animals presented a significant increase in the number of complex vocalizations (“modulated + complex” according to (Ey, et al. 2013)) which are the longer vocalization types emitted by mice and can last up to 0.15ms (Figure 3 B). Complex vocalizations have been recently associated to more social behaviours (Ey, et al. 2020). Therefore, in this experimental paradigm, designed to detect social vocalization, we observed a new partially-penetrant phenotype characterized by an increase in the number calls, and differences in vocal repertoire and “prosody” (Lahvis, et al. 2011).

**Figure 3:**
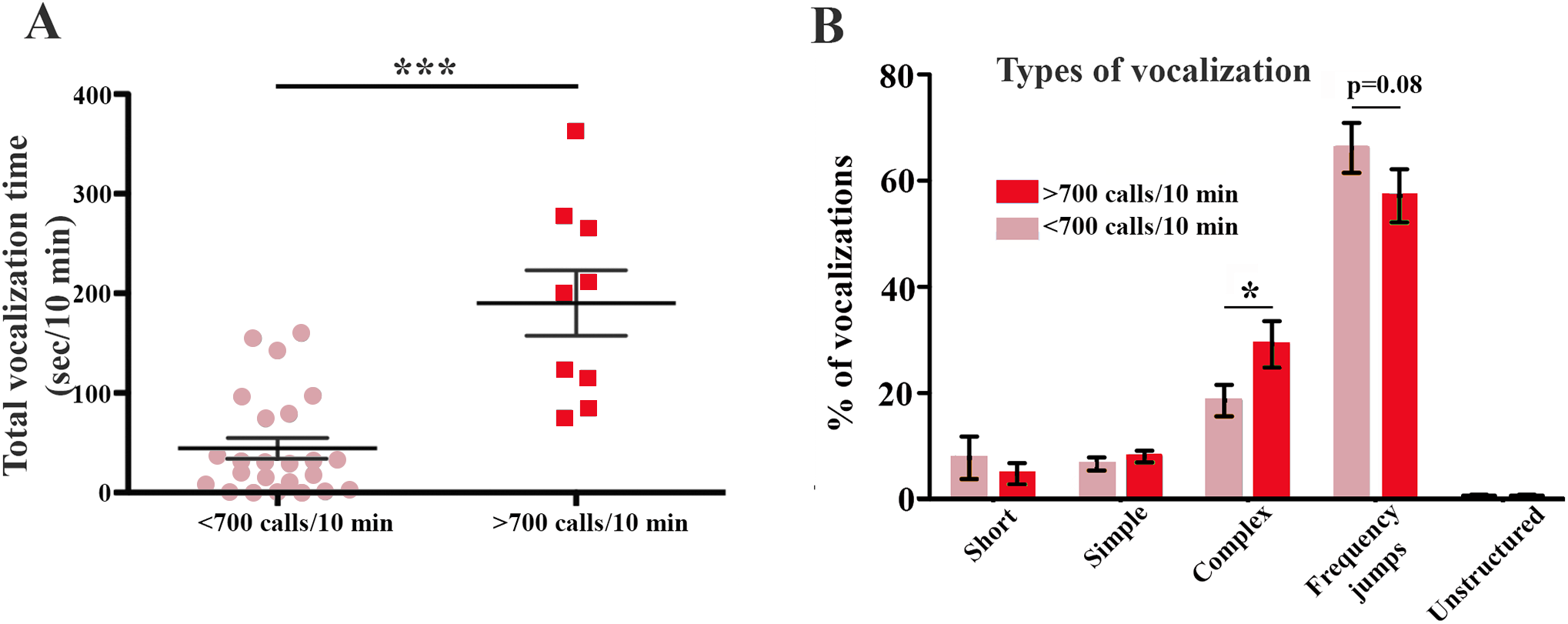
Total vocalization time and call types emitted by hyper-vocalizing and low-vocalizing mice. A) Total time spent vocalizing by mice emitting less or more than 700 calls/10 min. B) Percentage of call types produced by mice emitting less or more than 700 calls/10 min. Five types of emissions have been considered as in (Ey, et al. 2013): Short, Simple (Flat+Upward+Downward), Complex (Modulated + Complex), Unstructured, and Frequency jumps. Hyper-vocalizing animals vocalize, in average, four times more than low vocalizing animals and emit significantly more complex and modulated calls. (***= p<0.001; *=p<0.05, Mann-Whitney test).

### *Dlx5/6* expression in brain regions which control vocalization

Vocal communication depends on a complex neural network that includes regions of the forebrain, of the basal ganglia and of the brainstem (Jurgens 2002). These multiple inputs are integrated by the periaqueductal grey (PAG), a midbrain territory that is at the interface between the telencephalon and phonation/respiration motor centres. As all these brain territories include GABAergic neurons, we determined in which of them *Dlx5* is expressed. To this end we analysed the distribution of ß-galactosidase staining in *Dlx5^lacZ^* adult mouse brains (Figure 4) (Acampora, et al. 1999). We find that *Dlx5* is not expressed in the PAG or other midbrain regions (Fig. 4A) nor in posterior motor areas. On the contrary, *Dlx5* is strongly expressed in the cerebral cortex (Fig. 4B), in the hypothalamus (Fig. 4C) and in the striatum (Fig. 4D). These observations suggest that Dlx5/6 affect central cognitive and emotional inputs to vocal communication, but do not affect the “mechanics” of phonation.

**Figure 4:**
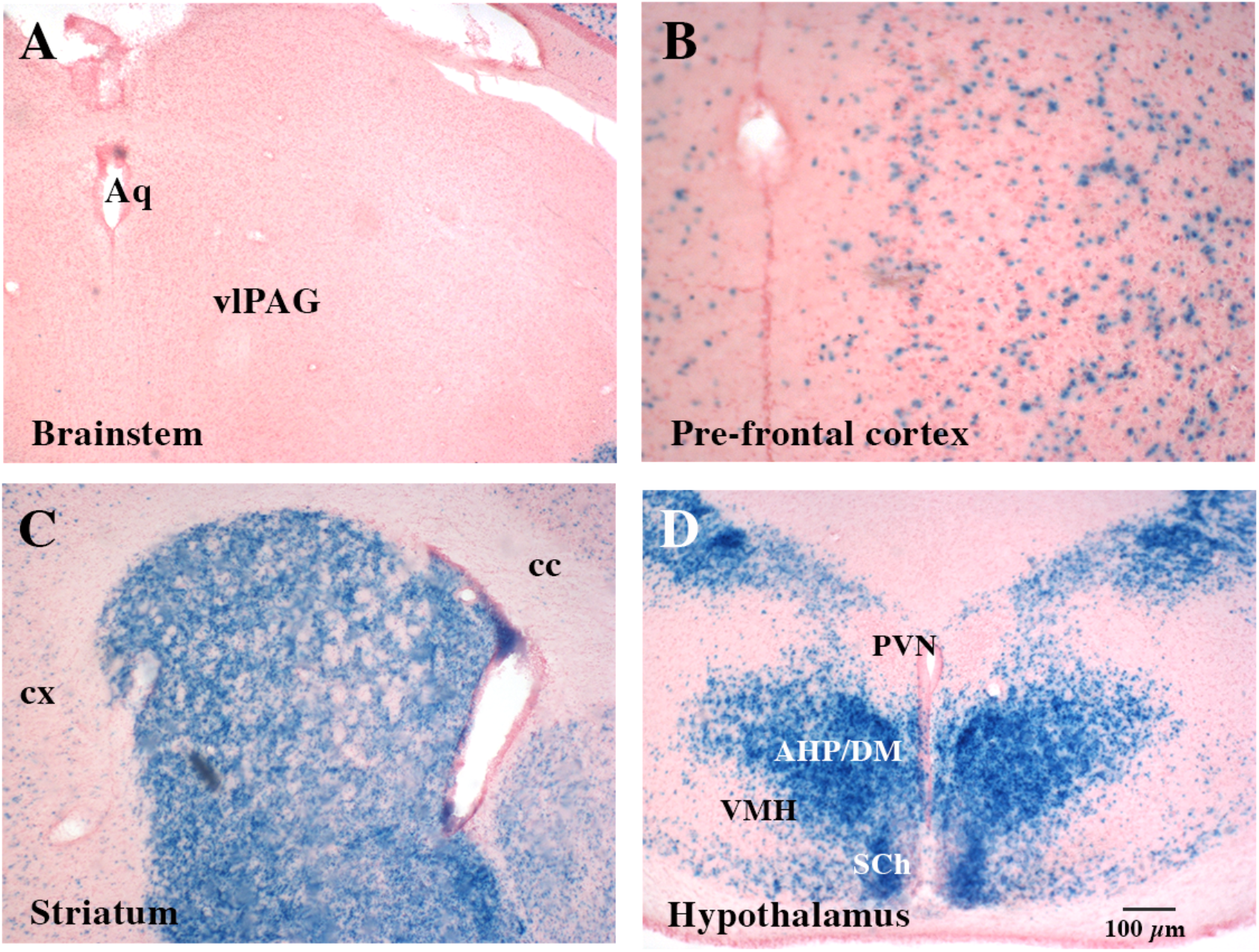
Expression of *Dlx5* in different adult brain regions involved in vocalization. X-gal staining on serial sections of *Dlx5^lacZ/+^* brains (Acampora, et al. 1999). *β*-D-galactosidase activity was not detected in the brainstem (A), in the cerebellum and in the midbrain including the ventro-lateral periacqueductal grey (vlPAG) a region which integrates sensory and cognitive inputs and controls vocalization motor centres. *Dlx5* was expressed in the prefrontal cortex (B) and in other cortical regions (cx), in the striatum (C) and in hypothalamic nuclei (D). AHP, Anterior hypothalamic area; cc, corpus callosum; cx, cortex; DM, Dorsomedial hypothalamus; PVN, para-ventricular nucleus; SCh, suprachiasmatic nucleus; VMH, ventro medial hypothalamus).

### Social behaviour of hyper-vocalizing mice

The social behaviour of the mice on which we had quantified UVs was analysed with the resident/intruder experimental setting described above. Behaviour and social interactions were filmed and analysed as previously described (de Chaumont, et al. 2019). Social behaviour between individuals of the same genotype was very variable resulting in no significant difference in sociability between control, *Dlx5/6^VgatCre/+^* and *Dlx5/6^VgatCre^* mice (de Lombares, et al. 2019). However, when we confronted the groups of hyper-vocalizing (>700 Calls/10min) and low-vocalizing (<700 calls/10min) animals, we found a highly-significant association between hyper-vocalization and hyper-sociability. Hyper-vocalizing animals spent more than twice time following or approaching the intruder than the low-vocalizing individuals (Figure 5 A, B).

**Figure 5:**
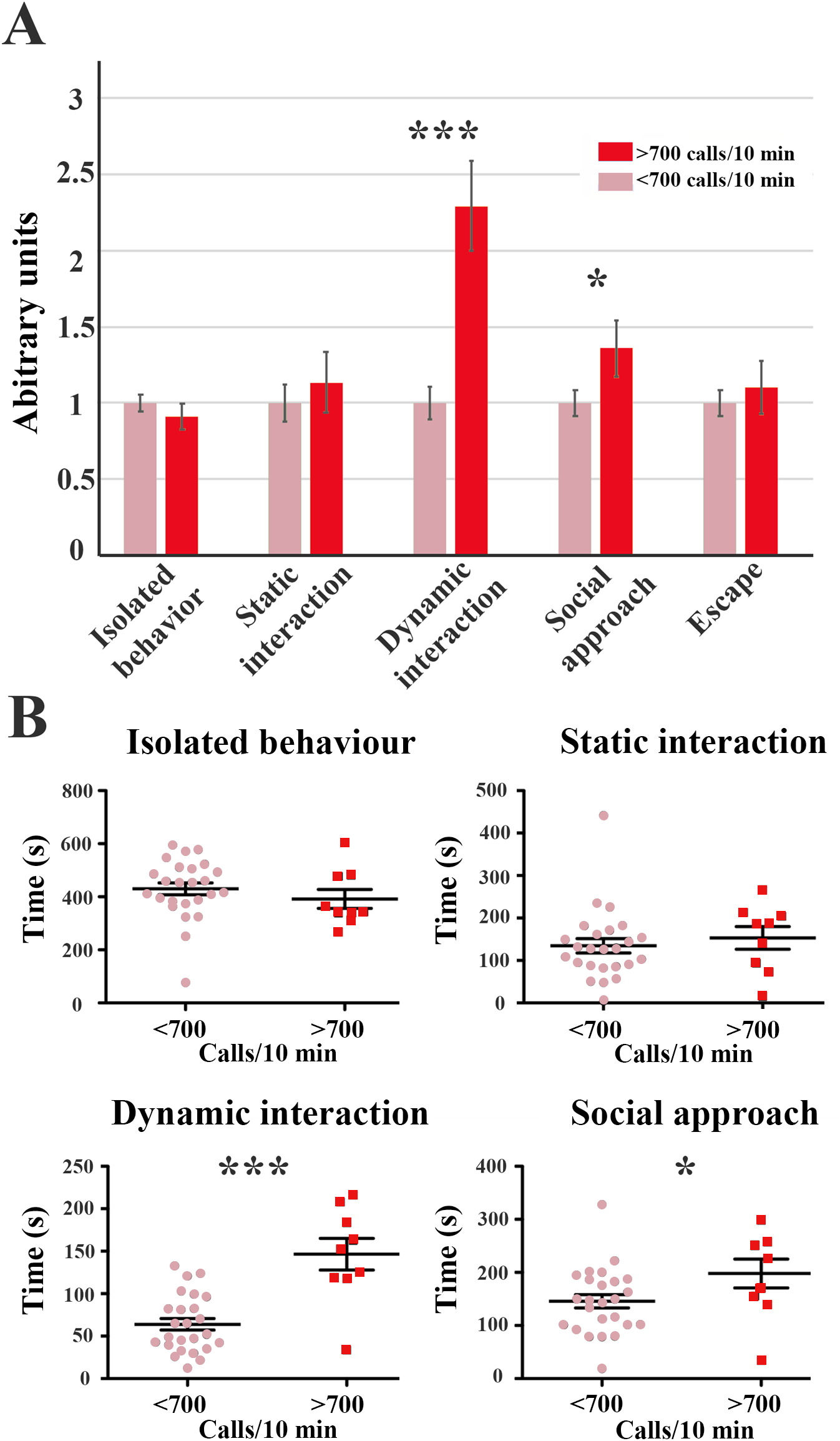
Social behaviour of hyper-vocalizing and low-vocalizing mice. The social interaction behaviour of *Dlx5/6^VgatCre/+^* (n=10), *Dlx5/6^VgatCre^* (n=9) and control (n=15) mice placed in the same cage with a control visitor was monitored using a protocol (de Chaumont, et al. 2019; de Lombares, et al. 2019) which associates computer vision and artificial intelligence to identify and categorize individual social interactions. The types of behaviour recorded were classified as follows: 1) Isolated behaviour (moving alone, rearing alone, stopped alone, jump alone, huddling alone), 2) Static interactions (rearing in contact, oral-oral contact, oro-genital contact, side-by-side in same or opposite orientation, 3) Dynamic interactions (moving in contact, train of 2 mice, following the same path), 4) Social approach (making contact, approaching in rearing, approaching), 5) Social escape (breaking contact, escaping), a detailed description of each behaviour is found in (de Chaumont, et al. 2019). The time spent by each mouse (sec) adopting a specific behaviour was measured over a 10 min recording. A) Normalized results: the average time spent in a specific behaviour by low-vocalizing animals was set as one (pink) and the ratio with the time spent in that same behaviour by hyper-vocalizing mice (red) is shown. For example, “Dynamic Interaction” represents a trait that was adopted 2.3 times more frequently by hyper-vocalizing mice than by low-vocalizing animals. B) Actual values in seconds recorded for each social behavioural trait. (***= p<0.001; *=p<0.05, Mann-Whitney test).

Therefore, the same animals in which the mutation of *Dlx5/6* in GABAergic neurons induced hyper-vocalization presented also a hyper-sociability phenotype.

### Introgression of a Neandertal-like haplotype in the *DLX5/6* locus

Patients carrying mutations in the *DLX5/6* locus or mouse models in which these genes are deregulated present abnormalities in at least five set of characters which differentiate *Homo sapiens* from *Homo Neanderthalensis:* 1) brain size and shape, 2) limb morphology, 3) craniofacial shape and 4) middle ear morphology and 5) bone density.

Comparing the *DLX5/6* locus of modern humans (Figure 6A) to those of Neanderthal (Altaï, Vindija), Denisovan, and other *Hominidae* we identified 298 SNPs with rare alleles that are shared between modern humans and Neanderthals and are absent from African populations in an interval of 400 kb centred on *Dlx5* coding sequences. Among them we identified, in the proximal region of *DLX5/6*, five SNPs (rs12704907, rs4729325, rs1005169, rs1724297, rs1207733), all located in non-coding regions, that co-segregate at more than 80% in individuals from the worldwide 1000 Genomes database (The 1000 Genomes Project Consortium), forming a “core” 37,160bp haplotype (*DLX5/6*-N-Haplotype) defined by all consecutive sites with Neandertal-like alleles (Table 1, Figure 6 A). This haplotype includes *DLX6* coding and regulatory sequences, the intergenic region between *DLX6* and *DLX5* and *DLX5* intron 3 and exon 3. Allele sharing with archaic genomes at polymorphic sites in non-African populations can result either from incomplete lineage sorting or admixture. To distinguish between these two scenarios, we looked at the length of the haplotypes. Introgressed haplotypes present fewer recombinations than those predicted from the time interval that separates modern humans from the common ancestral population. The *DLX5/6*-N-Haplotype is significantly longer than expected under the null hypothesis of incomplete lineage sorting (p value = 1.08e^−5^), supporting an archaic origin of the haplotype.

**Figure 6:**
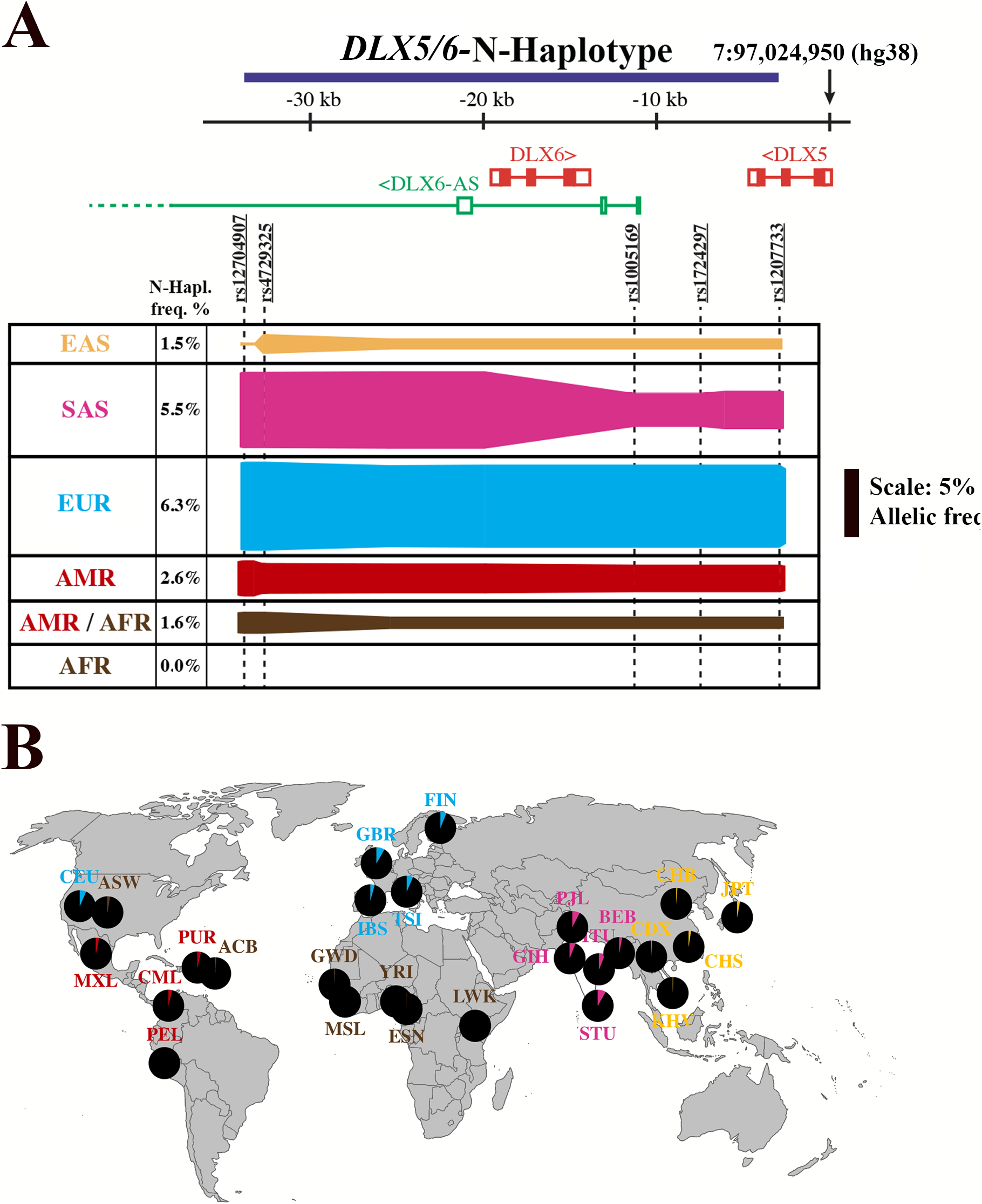
Linkage disequilibrium analysis of the distribution of the *DLX5/6-N*-Haplotype in different populations. A) For each population, the width of the colour bar represents the allele frequency for each SNP. SNPs linked by a solid bar cosegregate; if cosegregation is less than 100%, the width of the bar diminishes accordingly. The allele frequency of the most represented Neanderthal SNP of the *DLX5/6*-N-Haplotype is indicated for each population. The major SNPs constituting the haplotype are indicated. The scale bar represents a 5% allelic frequency within a population. EAS: East Asia populations; SAS: South Asia populations; EUR: European populations; AMR: American populations; AMR/AFR; Afro-americans; AFR: Africans. B) Geographical distribution of the *DLX5/6*-N-Haplotype. The allele frequency of the most represented SNP constituting the *DLX5/6*-N-Haplotype is represented on the earth map. The higher frequency of the *DLX5/6*-N-Haplotype is seen in Europe and South Asia, the haplotype is not present in African populations and in Peruvians from Lima. CHB: Han Chinese in Beijing, China; JPT: Japanese in Tokyo, Japan; CHS: Southern Han Chinese; CDX: Chinese Dai in Xishuangbanna, China; KHV: Kinh in Ho Chi Minh City, Vietnam; CEU: Utah Residents (CEPH) with Northern and Western European Ancestry; TSI: Tuscans in Italy; FIN: Finnish in Finland; GBR: British in England and Scotland; IBS: Iberian Population in Spain; YRI: Yoruba in Ibadan, Nigeria; LWK: Luhya in Webuye, Kenya; GWD: Gambian in Western Divisions in the Gambia; MSL: Mende in Sierra Leone; ESN: Esan in Nigeria; ASW: Americans of African Ancestry in South-West USA; ACB: African Caribbeans in Barbados; MXL: Mexican Ancestry from Los Angeles USA; PUR: Puerto Ricans from Puerto Rico; CLM: Colombians from Medellin, Colombia; PEL: Peruvians from Lima, Peru; GIH: Gujarati Indian from Houston, Texas; PJL: Punjabi from Lahore, Pakistan; BEB: Bengali from Bangladesh; STU: Sri Lankan Tamil from the UK; ITU: Indian Telugu from the UK

**Table 1:** SNPs included in the *DLX5/6*-N-Haplotype

| rs_ID | introgressed | Altai | Vindija | Denisova | Chimp | Bononbo | Gorilla | Orang |
| --- | --- | --- | --- | --- | --- | --- | --- | --- |
| rs12704907 | C | CC | CC | TT | T | T | T | T |
| rs4729325 | C | TT | CC | TT | T | T | T | T |
| rs1005169 | A | CC | CA | CC | C | C | C | C |
| rs1724297 | A | GG | AG | GG | G | G | G | G |
| rs1207733 | A | AA | AA | GG | G | G | G | G |

The proportion of individuals carrying the *DLX5/6*-N-Haplotype is variable between populations, from 1.5% in East Asia to 6.3% in Europe and absent in continental Africa (Figure 6A, B).

Overall, the proportion of the *DLX5/6*-N-Haplotype is low in East Asia and America (less than 4%) and completely absent from Africans and Peruvians from Lima. In South Asia and Europe, the *DLX5/6*-N-Haplotype is much more frequent reaching up to 8.2% in the British population of England and Scotland and in Sri Lankan Tamil in the UK. In African-American and Caribbean the haplotype is poorly represented (Figure 6B). The five reported SNPs are in LD in the European populations and the r2 are reported in Table 1S for different populations of the 1000 Genomes Project.

A haplotype (hg38 - chr7:96991129-97022094) of Neanderthal (Vindija) origin has been recently identified applying a two-state hidden Markov model to search for archaic fragments (Denisovan, Altai and Vindija Neanderthal) in the whole genome of the Icelandic population (see Supplementary Materials in (Skov, et al. 2020)). The identified haplotype corresponds to the *DLX5/6*-N-Haplotype. Although the *DLX5/6*-N-Haplotype is the most represented, some of the Icelanders present larger haplotypes which include several additional SNPs of Neanderthal origin. Similarly, in the 1000 Genomes database, the five “core SNPs”, present in every populations except Africa, cosegregate with other SNPs most of which correspond to those described in (Skov, et al. 2020) (Table 2S, Figure 1S).

### Association of the *DLX5/6* locus with human cognitive impairment and speech defects

Deletions or mutations at the *DLX5/6* locus are associated to Split Hand Foot Malformation Type 1 (SHFM1), a form of ectrodactyly in which limb malformations can be associated to cognitive, craniofacial or hearing defects. To evaluate the possible implications of the *DLX5/6* locus and of the *DLX5/6*-N-Haplotype in particular for human development and evolution, we reviewed the cognitive phenotypes reported for SHFM1-patients carrying known genetic alterations within this region (Rasmussen, et al. 2016). This analysis permitted to associate specific genomic sub-regions to syndromic traits such as mental retardation, language delay and autism (Figure 7).

**Figure 7:**
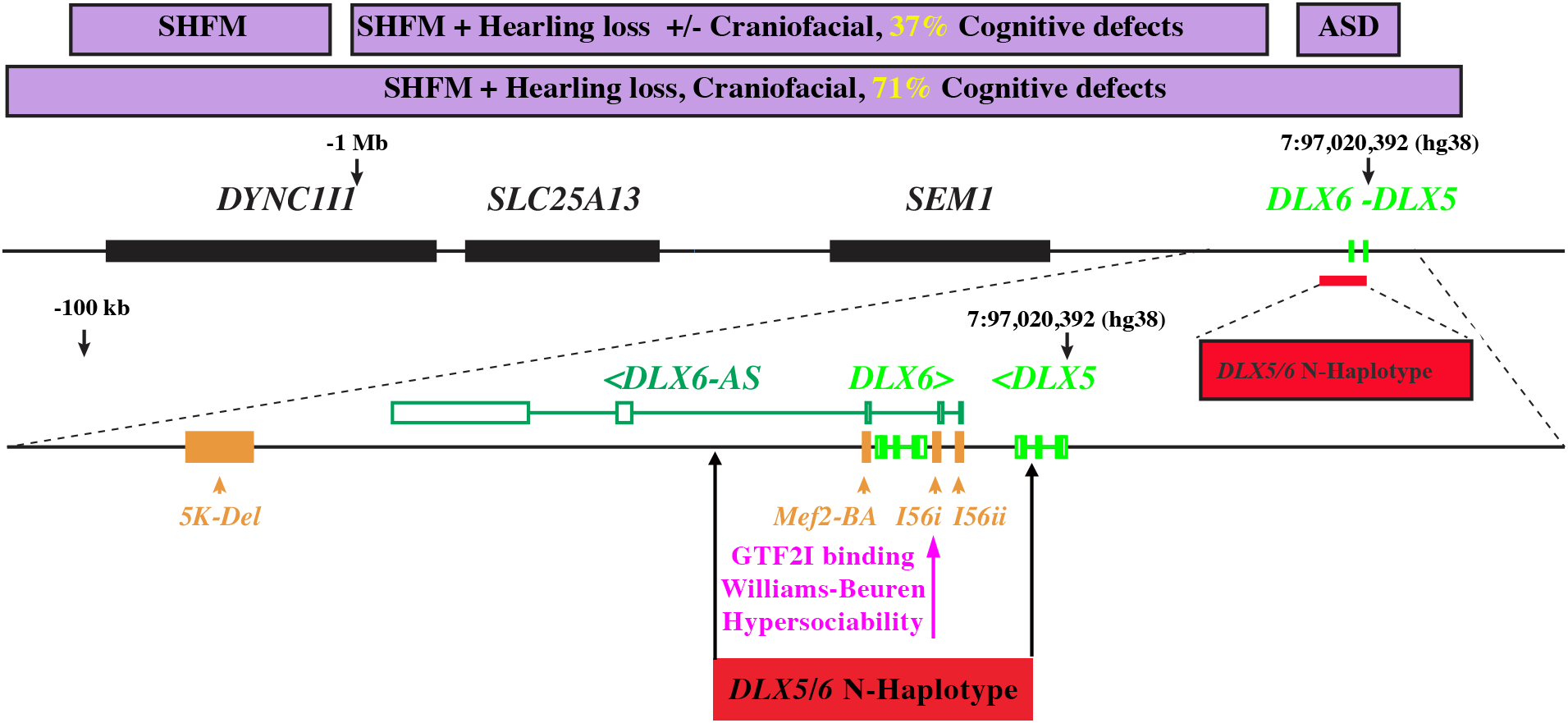
Association of different pathological conditions with the *DLX5/6* genomic region. The central black line represents the 7q21.3 locus, which includes at least 5 genes *DLX5, DLX6, SEM1, SLC25A13* and *DYNC1I1*. A long non-coding anti-sense RNA, *DLX6-AS*, is also transcribed from this region. Purple boxes in the upper part of the figure represent the phenotypic consequences observed in Split Hand Foot Malformation syndrome (SHFM) patients carrying deletion/mutations in this locus. While all SHFM patients present ectrodactyly, in certain cases this malformation is accompanied by craniofacial defects, hearing loss and/or cognitive defects. The closer the deletion are to *DLX5/6* coding sequences the higher is the frequency of patients presenting also cognitive impairment. Point mutations within the *DLX5/6* region and proximal or intragenic enhancers have been associated to autism spectrum disorders (ASD) without ectrodactyly or craniofacial defects and are included in the *DLX5/6*-N-Haplotype. Three conserved enhancers (*I56i, I56ii* and *Mef2-BA*) that direct *DLX5/6* expression in the brain (indicated in yellow) are also part of the haplotype. A point mutation within the *I56i* enhancer, prevents binding of GTF2I, a transcription factor deleted in Williams-Beuren syndrome, a condition characterized by hypervocalization and hypersociability (Poitras, et al. 2010).

Most families carrying mutations limited to the genomic region encompassing only limb-specific enhancers (Birnbaum, et al. 2012) present SHFM1 with no other physical or mental problems, with the exception of one case of mild cognitive defects. Families with deletions or translocations including craniofacial-specific enhancers present SHFM1 with hearing loss with or without other craniofacial anomalies and a high incidence of cognitive defects (Rasmussen, et al. 2016). Large heterozygous deletions covering both regulatory and coding *DLX5/6* sequences present SHFM1 with craniofacial defects associated in 5 out of 7 cases to severe mental retardation. Patients carrying mutations restricted to *DLX5/6* intergenic region or their coding sequences have a higher probability to include members with ASDs or speech delay but do not present the limb deformities of SHFM1 (Hamilton, et al. 2005; Nakashima, et al. 2010; Poitras, et al. 2010; Chang, et al. 2011).

The *DLX5/6*-N-Haplotype includes enhancers (*I56i, I56ii* and *MEF2*) capable to direct *DLX5/6* expression in the brain (Fazel Darbandi, et al. 2016; Assali, et al. 2019). Interestingly a polymorphism in enhancer *I56i* is associated to the binding of GTF2I, a transcription factor deleted in Williams-Beuren syndrome, a complex condition characterized by craniofacial, hyper-vocalization and hyper-sociability (Figure 7) signs which has been considered an entry point to understand the evolution of the modern human face and prosociability (Zanella, et al. 2019).

Through PheWeb we examined the association between the SNPs constituting the *DLX5/6*-N-Haplotype and the phenomes of the Michigan genomics initiative and of the UK Biobank. Results with p-values < 0.01 are reported in Supplementary Materials. Table 3S shows phenotypes identified in the UK Biobank dataset through a generalized mixed model association test that uses the saddlepoint approximation to account for case-control imbalance (SIAGE) and adjusting for genetic relatedness, sex, birth year and the first 4 principal components. Table 4S indicates phenotypes identified in the UK Biobank dataset using a least-squares linear model predicting the phenotype with an additive genotype coding (0, 1, or 2 copies of the minor allele), with sex and the first 10 principal components from the UK Biobank sample QC file as covariates. Table 5S reported the phenotypes identified in the Michigan Genomics.

The phenotype “*Cause of death: other specified respiratory disorders*” (Table 4S and Table 6S) has the highest positive association to the introgressed variant of alleles rs4729325 (C-allele) and rs 12704907 (C-allele) (p-value 2*10^−5^). ClinVar, GWAS Catalog, Gwas Central, GTEX PORTAL, clinGen were examined for the same SNPs but no results were found.

The GeneATLAS analysis showed that the most significant phenotypes are those associated to adipose tissue accumulation and “Standing height”. A detailed list of all phenotype associated to each SNPs is reported in Supplementary Material (from Table 7S to Table 11S).

### The *DLX5/6*-N-Haplotype allele is under-represented in semi-supercentenarians

As the *DLX5/6* locus has been associated to ASDs and mental retardation ((Gandal, et al. 2018) and (Figure 7)), we analysed the *DLX5/6*-N-Haplotype frequency in the MSSNG database of ASDs families (RK, et al. 2017) and we found a non-significant association (p=0.16) with the disease. This finding, however, cannot exclude an association of the haplotype to specific mental traits not necessarily represented in the ASDs database. We then focused on other traits that can be associated to social vocalization. Notably, mortality risk has been correlated to social environment in humans and other social mammals (Snyder-Mackler, et al. 2020). Since, inactivation of *Dlx5/6* in GABAergic neurons results in a 33% life span extension in mice (de Lombares, et al. 2019) we then focused on the population of semi-supercentenarians (CEN), i.e. people over 105 years of age (105+) characterized by exceptional longevity. The frequency of the *DLX5/6-N*-Haplotype or of truncated haplotypes containing at least 2 consecutive Neandertal SNPs were calculated for all the European populations present in the 1000 Genomes database and estimated for a cohort of 81 Italian semi-supercentenarians. Fixation index (F_st_) analysis showed a significant difference in *DLX5/6*-N-Haplotype frequency (p-value < 0.05). The 105+ (CEN) presented a 50% under-representation of the haplotype when compared to any other European population with the only exception of IBS and constituted the most divergent group (Figure 8A).

**Figure 8:**
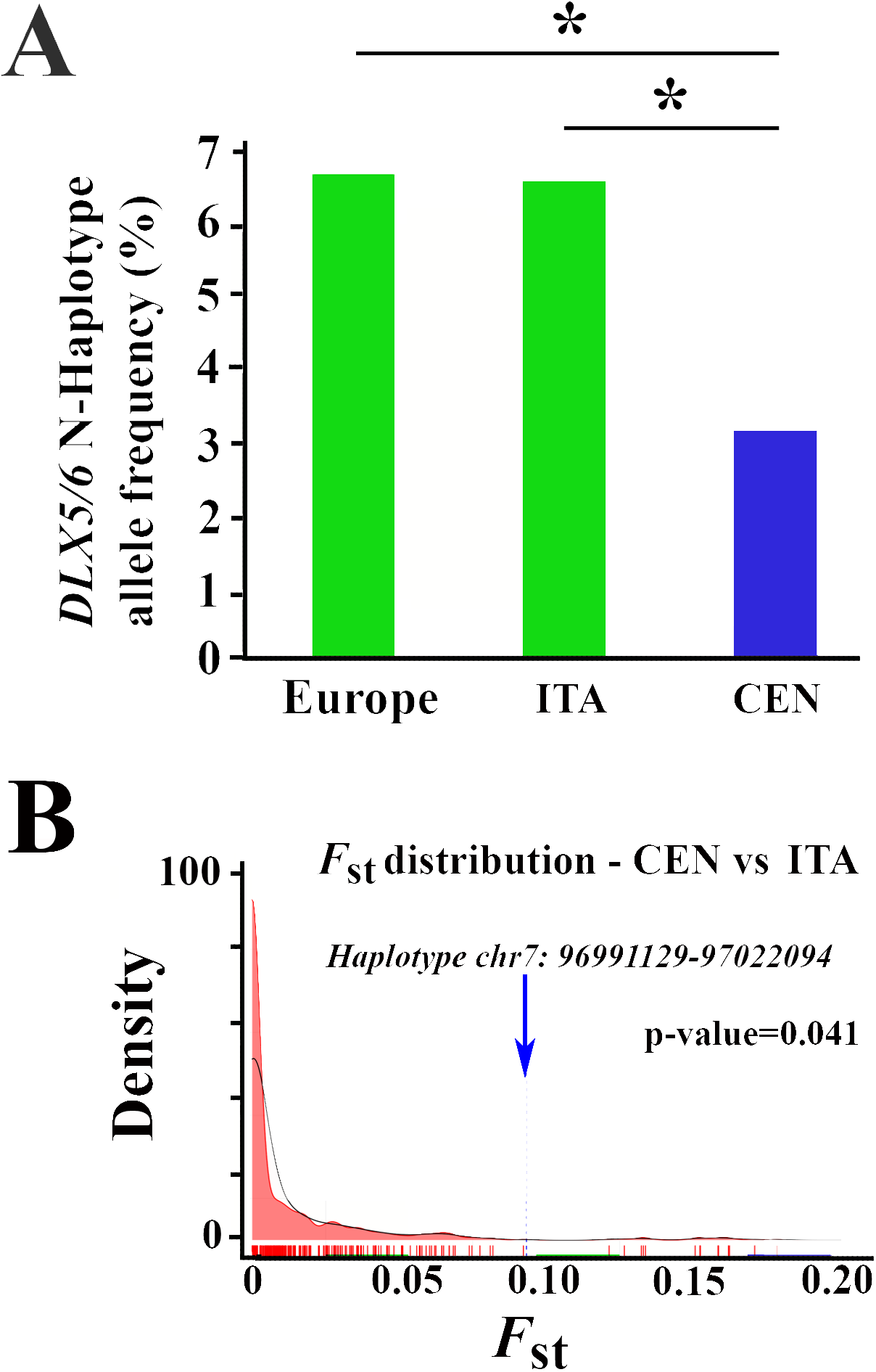
Allele frequency of the *DLX5/6*-N-Haplotype in the semi-supercentenarian population. A) The allele frequency of the archaic form of the *DLX5/6*-N-Haplotype of the European populations present in the 1000 Genomes Project is confronted to the CEN population of 81 semi-supercentenarians (105+ yo) and to the ITA population obtained combining the Tuscany 1000 Genomes population (TSI) with 36 controls genomes matching the different geographical origin of CEN. While the ITA population does not show any significant difference from the Europeans a significant reduction of the *DLX5/6*-N-Haplotype allele frequency is observed in the semi-supercentenarian population (*: p<0.05, *F*_st_ distribution). B) Density plot build considering all the *F*_st_ values calculated considering more than 600 archaic haplotypes (described in (Skov, et al. 2020)) for CEN vs ITA groups. The dashed line indicates the value of *F*_st_ calculated for *DLX5/6*-N-Haplotype.

Then we performed a comparison between CEN and the Italian control group termed “ITA” (the group closest to CEN in term of genetic ancestry and composed by TSI from 1000 Genomes Project and 36 Italian healthy individuals from the Italian peninsula) also showed a significant difference (p-value = 0.026). Chi-square analysis based on the allelic distribution between CEN and the total European population also showed significant underrepresentation of the haplotype in CEN (p<0.05) (Figure 8A).

Since the possibility of bias due to small sample size we also calculated *F*_st_ between CEN, ITA and European populations from 1000 Genomes Project (TSI, GBR, FIN, IBS, CEU), selecting a list of more than 600 archaic haplotypes located in chromosome 7 from the recent paper of (Skov, et al. 2020). We performed a distribution of *F*_st_ computed between all population pairs (CEN, ITA and TSI, GBR, FIN, IBS, CEU of 1000 Genomes Project) and for each distribution we checked if the *F*_st_ calculated for *DLX5/6*-N-Haplotype is significantly different from the reported distribution. The comparisons of CEN with the Italian control group termed (ITA) showed a significant difference (p-value = 0.041). The *F*_st_ of *DLX5/6-N*-Haplotype in other comparisons including CEN (CEN vs FIN, CEN vs CEU, CEN vs GBR and CEN vs TSI) is always greater than the 90th percentile of the empirical distribution with the only exception of CEN vs IBS. *Fst* distribution of all archaic haplotypes for CEN vs ITA is reported in Figure 8B and the *Fst* value related to *DLX5/6*-N-Haplotype is indicated with a blue dashed line, suggesting that the selected haplotype significantly differentiates the two populations (p-value=0.041).

## DISCUSSION

Here we have shown that reduction of *DLX5/6* expression in mouse GABAergic neurons results in a hyper-vocalization and hyper-socialization phenotype. Furthermore, we identify a *DLX5/6*-Neanderthal-derived haplotype present in up to 8% of non-African alleles that includes enhancers controlling gene expression in the central nervous system (Fazel Darbandi, et al. 2016) and sequences associated to either hyper-or hypo-socialization/vocalization pathologies such as Williams-Buren syndrome (Poitras, et al. 2010) or ASDs (Gandal, et al. 2018). Interestingly the *DLX5/6*-N-Haplotype is underrepresented in the genomes of healthy semi-supercentenarians suggesting an association between *DLX5/6* brain expression and healthy aging reminiscent of what has been shown in mouse models (de Lombares, et al. 2019).

In social mammals, such as mice and humans, vocal communication (VC) is essential for the survival of the group. VC is needed to convey diverse information such as location, social status, reproductive availability, emotional context (fear, aggression), food availability and presence of predators. Such a vast set of inputs and outputs is coordinated by a complex neural network that combines regions of the forebrain, of the basal ganglia and of the brainstem (Jurgens 2002). A key brain region involved in the integration of these neuronal activities is the periaqueductal grey (PAG), a midbrain territory that receives multiple telencephalic inputs and transmit them to phonation and respiration motor centres. As *Dlx5/6* are not expressed in the PAG, but only in telencephalic interneurons, their effect on vocalization must derive from cognitive initiation and modulation and not from impairment of laryngeal or respiratory motor centres.

In comparison to their normal controls, many *Dlx5/6^VgatCre^* and *Dlx5/6^VgaCre/+^* mice emit a higher number of longer “complex” calls often not separated by any interruption. These long-modulated USVs are emitted prevalently when mice are engaged in socially-intense behaviours and more rarely during isolated behaviour (Ey, et al. 2020). Therefore, the high-vocalization profile of *Dlx5/6^VgatCre^* mice is predictive of an enhanced social behaviour and is in line with our finding of an increase in dynamic socialization of high-vocalizing individuals.

Many mouse models have been generated to study the effect of genetic lesions associated to human neurodevelopmental disorders. In particular, the simultaneous study of ultrasonic vocalizations and behaviour has permitted to evaluate the impact of specific mutations on social communication and interaction in mouse models of ASDs (Caruso, et al. 2020). For example, female-female social interactions and USV have been analysed in *Shank2^−/−^* and *Shank3^−/−^* mutant mice showing reduction in the number and duration of USVs associated to diminished sociability (Schmeisser, et al. 2012; Ey, et al. 2013; Wohr 2014). Extrapolation of our results to the human condition would suggest that reduced *DLX5/6* expression in GABAergic neurons could be associated to increased social vocalization, a phenotype opposite to ASDs.

Our understanding of the origin of psychiatric pathologies such as schizophrenia and ASDs has recently progressed substantially thanks to large Genome Wide Association studies (GWAs) which have shown that, beyond some specific cases where monogenic mutations cause highly penetrant phenotypes, psychiatric conditions are complex polygenic disorders associated with hundreds of common genetic loci involving genes associated to synaptic function and neural development (Huguet, et al. 2016; Dennison, et al. 2019). Module analysis of gene-networks implicated in interneuron-dependent ASD and schizophrenia has identified gene expression variations and splicing of *DLX5*, *DLX6* and *asDLX6nc*. Indeed, the modules including these *DLX* genes and other key genetic components of GABAergic neurons constitute the major hub for both ASD and schizophrenia (Gandal, et al. 2018). Remarkably, a point mutation within the *I56i* enhancer, is associated with binding of the transcription factor GTF2I which is deleted in Williams-Beuren syndrome characterized by hypervocalization and hypersociability (Poitras, et al. 2010).

*GTF2I* is located in 7q11.23, a genomic region which deletion or microduplication provokes complex syndromes with opposite social and speech language behaviours including sociability, speech language and cognitive alteration associated with neural crest-associated facial features (López-Tobón, et al. 2020). These genetic networks, which include *DLX5/6* and control concomitant modifications of craniofacial morphology, cognition and sociability are proposed to have contributed to human self-domestication (Zanella, et al. 2019). Furthermore, *DLX5/6* were proposed as potential contributors to brain and skull globularization and acquisition of human-specific language capacities (Boeckx and Benitez-Burraco 2014a).

The origin of vocal language in the genus *Homo* has been approached through anatomical analyses of the brain and larynx and, more recently, through the comparison of genes involved in language acquisition such as *FOXP2* (Dediu and Levinson 2013; Dediu and Christiansen 2016). Remarkably *FOXP2* is located in 7q31, close to the *DLX5/6* locus (7q21), and regulates the expression of a large antisense noncoding RNA of *DLX6 (asDLX6nc* or *Evf1/2*) (Vernes, et al. 2006; Vernes, et al. 2011), which is directly involved in the control of *Dlx5* and *Dlx6* expression (Feng, et al. 2006).

The *DLX5/6*-N-Haplotype covers the complete sequence of *DLX6*, the intragenic region and the 2^d^ intron and 3^rd^ exon of *DLX5* and three known enhancers for brain expression of *DLX5/6*: *I56i, I56ii* and *Mef2-BA* (Birnbaum, et al. 2012), and the three first exons of the long non-coding RNA *asDLX6nc*, which has been implicated in GABAergic neuronal differentiation (Cajigas, et al. 2018). The major SNPs composing the *DLX5/6*-N-Haplotype are located in non-coding sequences in proximity of the known enhancers and splice sites of *DLX5* and *asDLX6nc* and could well alter their expression profile.

Whole genome analyses of Neanderthal and Denisovan introgressions in 27,566 Icelanders identify the presence of *DLX5/6*-N-Haplotype comprising the 5 core SNPs with a 1.8% allelic frequency identified as having a Neanderthal origin. This introgression coexists with other, rare, longer haplotypes (Skov, et al. 2020).

The comparison of human genomes with those of other vertebrate species and, in particular, with other *Hominidae*, has revealed the presence of Human Accelerated Regions (HARs), short sequences presenting an important number of Single Nucleotide Changes (SNCs) specific to *Homo sapiens* (Lindblad-Toh, et al. 2011; Capra, et al. 2013). Interestingly, the *DLX5/6* locus includes at the same time 5 HARs and, as shown in this report, an introgressed Neanderthal region.

This paradoxical situation might suggest that enhancers implicated in non-neural functions of *DLX5/6* such as bone remodelling and reproduction could have been strongly counter-selected, whereas brain-expression driving enhancers could have been conserved in the human population. Interestingly, two of these HARS are located immediately in 5’ from *DLX5/6*-N-Haplotype (2.1 and 3.1 kb), whereas the other three are more distant (81.5, 634, 1088 kb upstream although in the range of the multiple known modifiers of *DLX5/6* expression). The presence of the proximal HARs may have limited the extension of the introgressed haplotype in 5’.

Vocal communication and socialization are pivotal traits of the evolution of modern human society. The conservation of the *DLX5/6*-N-Haplotype in 6.3% of EUR alleles (which represent about 12,6% of the population as the *DLX5/6*-N-Haplotype is almost invariably heterozygous), suggests that its presence in a proportion of individuals is beneficial for the sustainability of modern human society. As *DLX5/6* are pleiotropic genes with multiple interconnected functions including craniofacial morphogenesis, neuronal differentiation, bone development and reproduction (Merlo, et al. 2000) one could speculate that the presence of the *DLX5/6*-N-Haplotype might also affect other traits characteristic of the evolution of *Homo sapiens*, such as metabolism, skull shape and bone density. In particular one of the main differences between modern humans and Neanderthals has been the composition and size of social groups: while Neanderthal groups were small, composed by 10 to 30 individuals, modern humans developed complex societies (Duveau, et al. 2019). It is possible, therefore, that the *DLX5/6*-N-Haplotype could influence characters affecting the structure of modern human society.

However, the *DLX5/6*-N-Haplotype seems to correlate with different aspects of human health in modern human populations and some of them are pathological conditions. This observation is in line with data that demonstrates that high proportion of Neanderthal introgression might be deleterious for modern humans (Juric, et al. 2016). Thus, we can speculate that the frequency of this haplotype in modern human population was the result of balancing selection shaped by different evolutionary forces. Future studies are needed to address this point.

Interestingly the *DLX5/6*-N-Haplotype is underrepresented in semi-supercentenarians, a population considered as a paradigm of healthy ageing as it has the capacity to slow the progression of pathological conditions determining reduced mental and physical functions (Vacante, et al. 2012).

This observation could be related to the 33% lifespan extension described for *Dlx5/6^VgatCre^* mice, suggesting that the *DLX5/6*-N-Haplotype could regulate *DLX5/6* neuronal expression (de Lombares, et al. 2019). These data on semi-supercentenarians could be seen in the light of George Williams antagonistic pleiotropy theory (Williams 1957) who proposes that genes controlling more than one phenotypic trait can have been positively selected being favourable early in life and in adulthood but can become detrimental later in life (old age and extreme longevity), phases of life subjected to weaker evolutionary pressures (Williams 1957; De Benedictis and Franceschi 2006; Giuliani, et al. 2017).

Our findings could well be integrated within current ideas on the biology of ageing which suggest that ageing rates, and consequently lifespans, evolve as a function of trade-offs with other characters including social, cognitive and reproductive capacities (Yang, et al. 2016; Giuliani, et al. 2018; Snyder-Mackler, et al. 2020).

## MATERIALS AND METHODS

### Animals

Procedures involving animals were conducted in accordance with European Community (Council Directive 86/609) and French Agriculture Ministry directives (Council Directive 87–848, permissions 00782 to GL). The project was reviewed and approved by the “Cuvier” ethical committee of the Muséum National d’Histoire Naturelle (approval n° 68-028r1) validated by the French Ministry of Agriculture.

Mice were housed in light, temperature and humidity-controlled conditions. Food and water were available ad libitum. Mice were individually identified by microchip implanted 3 weeks postnatal. Litter sizes and genotypes were kept in record. *Dlx5/6^VgatCre/+^* and *Dlx5/6^VgatCre^* mice, were generated using a Cre/*lox* strategy to simultaneously remove *Dlx5/6* homeodomain-coding sequences specifically in GABAergic neurons (de Lombares, et al. 2019): *Dlx5/6^flox/flox^*mice (Bellessort, et al. 2016) were crossed with *Slc32a1^tm2(cre)Lowl^* knock-in mice (here referred as *Vgat^cre/+^* mice) in which the “*Cre-recomhinase*” gene is expressed only by GABAergic neurons on a mixed C57BL6/N X DBA/2 genetic background. Phenotypes of *Dlx5/6^VgatCre/+^* and *Dlx5/6^VgatCre^* mice were compared to those of control littermates.

### Vocalization and social behaviour analysis

Test females were isolated 3 days before the trial to increase their motivation for social interactions. On trial day, the tested female mouse was placed in a test cage (Plexiglas, 50 cm × 25 cm × 30 cm, 100 lux, with fresh bedding) for 30 min habituation. After this time, an unfamiliar group-housed control adult female was introduced. Ultrasonic vocalizations were recorded with a condenser ultrasound microphone Polaroid/CMPA, the interface UltraSoundGate 416-200 and the software Avisoft-SASLab Pro Recorded from Avisoft Bioacoustics (sampling frequency: 300 kHz; FFT-length: 1024 points; 16-bit format). Ultrasonic vocalizations were recorded for the pair of mice tested (one control newcomer and an habituated control, *Dlx5/6^VgatCre/+^* or *Dlx5/6^VgatCre^* mouse). All analysed animals were 6 months old. Previous studies suggest that the contribution of the newcomer to the vocalization is minor in comparison with that of the occupant (Hammerschmidt, et al. 2012). We measured the latency for the first ultrasonic vocalizations and the total number of ultrasonic vocalizations emitted. We manually established the distribution of ultrasonic vocalizations among the following call types (see (Ey, et al. 2013)): 1) “*short*”, call duration less than 5 ms; 2) “*simple*”, call duration longer than 5 ms and a frequency range either smaller than 6.25 kHz (*simple flat*) or with a frequency modulation in only one direction (*simple upward* or *downward);* 3) “*complex*”, frequency modulations in more than one direction and frequency range larger than 6.25 kHz (*modulated*), or inclusion of one or more additional frequency components (harmonic or non-linear phenomena, but no saturation); 4) “*unstructured*”, no pure tone component and 5) “*frequency jumps*”, presence of one or more jump(s) in frequency, without time gap between the consecutive elements.

We used unpaired non-parametric Wilcoxon tests to compare the latency for the first call and the call rate between genotypes. We used Chi-squared (with a Bonferroni correction for multiple testing) tests to compare the proportions of calls between genotypes.

During vocalization analysis, animals were filmed for 10 min. Social behaviour was measured using a real-time approach that couples computer vision, machine learning and Triggered-RFID identification to track and monitor animals (de Chaumont, et al. 2019).

### Histological analysis

Mice were fixed by transcardiac perfusion with 4% paraformaldehyde and brains were post-fixed by overnight immersion in 4% paraformaldehyde at 4°C. Samples were cryoprotected in 30% sucrose and frozen. Cryoprotected brains were embedded in OCT (Leica, France) and 60-μm-thick free-floating cryostat sections were prepared.

For *lacZ* expression analysis, X-gal staining was performed as described (Acampora, et al. 1999) on sections of *Dlx5^lacZ/+^* brains. Pictures were acquired using a Leica SP5 confocal microscope.

### Identification of *DLX5/6*-Neanderthal haplotype introgression

To investigate whether polymorphisms identified in the *DLX5/6* locus belong to a Neanderthal introgressed haplotype, we analysed individuals from the 1000 Genomes dataset (The 1000 Genomes Project Consortium phase 3, 2015) and looked for Neandertal-like alleles in a window of 200kb up and downstream from *DLX5*, using continental African populations as outgroup. Among these SNPs, we identified an haplotype of 37,160bp (GRCh 38 – 7: 96,991,129 – 97,022,094) defined by five consecutive SNPs (rs12704907, rs4729325, rs1005169, rs1724297, rs1207733) with Neandertal-like alleles shared by at least 80% of individuals worldwide. We computed the probability of generating such a long haplotype under a scenario of incomplete lineage sorting by assuming the haplotype length follows a gamma distribution (Huerta-Sanchez, et al. 2014) using the parameters from (Dannemann, et al. 2016) and the local recombination rate of 2.046 cM/Mb (Hinch, et al. 2011). The haplotype is significantly longer than expected under the null hypothesis of incomplete lineage sorting (p value = 1.08e^−05^), supporting an Archaic origin. The *DLX5/6*-N-Haplotype covers part of *DLX5*, all *DLX6*, the intergenic region and regulatory region upstream of *DLX6*. The proportion of each SNP in the different geographic regions was derived from the 1000 Genomes database phase 3, which included 694 American, 1008 East-Asian, 1006 European, 978 South-Asian and 1322 African genomes.

### *DLX5/6*-N-Haplotype allelic frequency

The allelic frequency of each SNP composing the *DLX5/6*-N-Haplotype and their linkage disequilibrium was determined in populations of different geographical and ethnical origins present in the 1000 Genomes dataset (The 1000 Genomes Project Consortium, 2015). These frequencies were used to reconstruct the haplotype using the linkage disequilibrium R2 between the different SNPs, their variations suggested the coexistence of several haplotypes of different size (unphased reconstruction). Genome-wide phased data (Skov, et al. 2020) also identified the introgression of the *DLX5/6*-N-Haplotype and the other haplotypes of different size from Neanderthal.

*DLX5/6*-N-Haplotype representation in autistic populations was determined using the MSSNG database (RK, et al. 2017) and the probability was calculated by transmission disequilibrium test (TDT).

PheWeb was examined to identify phenotypes associated to the five SNPs (rs12704907, rs4729325, rs1005169, rs1724297, rs1207733) (Gagliano Taliun, et al. 2020). We examined UK Biobank dataset through PheWeb based on SAIGE analysis of ICD - derived traits (http://pheweb.sph.umich.edu/SAIGE-UKB/) (Zhou, et al. 2018) and on the first Neale lab analysis (https://pheweb.org/UKB-Neale/). PheWeb was examined to identify association considering participants of the Michigan Genomics Initiative (http://pheweb.org/MGI-freeze2/).

PhenoScanner v2 (http://www.phenoscanner.medschl.cam.ac.uk/), ClinVar (https://www.ncbi.nlm.nih.gov/clinvar/), GWAS Catalog (https://www.ebi.ac.uk/gwas/), Gwas Central (https://www.gwascentral.org/), GTEX Portal (https://gtexportal.org/home/), clinGen (https://clinicalgenome.org/) were examined to identify in silico potential functional impact of the five SNPs (rs12704907, rs4729325, rs1005169, rs1724297, rs1207733). Then we analysed phenotypic associations in the UK Biobank using the GeneATLAS tool (Canela-Xandri, et al. 2018). In brief, the Gene ATLAS provides phenotypic associations based on Mixed Linear Models using 452,264 Britons with European descent. The model includes sex, array batch, UK Biobank Assessment Centre, age, and 20 genomic principal components. Results are reported in Supplementary Materials (Table 7S-11S).

The genomes from a cohort of 81 healthy semi-supercentenarians and supercentenarians [105+/110+] older than 105 years (mean age 106.6 ± 1.6) recruited in North, Centre and South of the Italian peninsula (CEN) were compared to a population (ITA) obtained combining the Tuscany 1000 Genomes population (TSI) with 36 controls genomes matching the different geographical origin of CEN. European population correspond to TSI, CEU, FIN, GBR and IBS people present in 1000 Genomes. Haplotypes were used to compute the F_st_ statistic (Wright 1951) between all the combinations of population pairs. To evaluate the significance of the F_st_ observed values we performed 500 simulations, where individuals were randomly assigned to one of the seven studied populations, and for each simulation F_st_ statistic was computed. At the end of this procedure we were able to fit a probability density function for the F_st_ statistic and to compute an empirical p-value for the F_st_ using the *GHap* (Utsunomiya, et al. 2016) R package. The significance of allelic differences between European population and semi-supercentenarians was also determined with a Chi-square test.

In order to test if the *F*_st_ calculated for the *DLX5/6*-N-Haplotype is within the distribution of other archaic haplotypes or is an outlier, we selected 620 archaic haplotypes (Skov, et al. 2020) located on the same chromosome (chr7) and phased each of the 620 individual genomes by applying SHAPEIT software v.2.17 (Delaneau, et al. 2013; O’Connell, et al. 2014). For each archaic haplotype, the *F*_st_ statistics (Wright 1951) was computed between all population pairs (CEN, ITA and CEU, FIN, GBR, IBS from 1000 Genomes Project). Then, for each pairwise comparison, we fitted a probability density function for the *Fst* statistic and an empirical p-value was computed for the *Fst* value calculated on our haplotype. The *Fst* analysis was implemented in R software (R Core Team, 2019) using the GHap package (Utsunomiya, et al. 2016).

## Supporting information

Supplementary_Levi

## ACKNOWLEDGMENTS

This research was partially supported the ANR grants TARGETBONE (ANR-17-CE14-0024) and METACOGNITION (ANR-17-CE37-0007) to GL, grants of the “Fondation-NRJ – Institut de France” (n° 216612) and the ATM “Cognitio” to NNN, CdL is supported by a grant of the French Ministry of Research.

We thank the following researchers for their valuable input:

Dr. Stéphane Peyregne, Department of Evolutionary Genetics, Max Planck Institute for Evolutionary Anthropology, Leipzig, Germany for analysis of the *DLX5/6*-N-Haplotype.

Dr.s Thomas Bourgeron, Elodie Ey and Fabrice de Chaumont from the Pasteur Institute, Paris, for initial analysis of the vocalization and behavioural data.

Dr Robbie Davies from The Centre for Applied Genomics, Toronto, Canada for analysis of the association between the *DLX5/6*-N-Haplotype and autism in the MSSNG database.

A particular thank goes to the team in charge of mouse animal care and in particular M Stéphane Sosinsky and M Fabien Uridat and to Pr. Amaury de Luze in charge of the “Cuvier” ethical committee. We thank Mss. Aicha Bennana and Lanto Courcelaud for administrative assistance.

